# Concordance of Genetic Variation that Increases Risk for Tourette Syndrome and that Influences its Underlying Neurocircuitry

**DOI:** 10.1101/366294

**Authors:** Mary Mufford, Josh Cheung, Neda Jahanshad, Celia van der Merwe, Linda Ding, Nynke Groenewold, Nastassja Koen, Emile R. Chimusa, Shareefa Dalvie, Raj Ramesar, Psychiatric Genomics Consortium - Tourette Syndrome working group, James A. Knowles, Christine Lochner, Derrek P. Hibar, Peristera Paschou, Odile A. van den Heuvel, Sarah E. Medland, Jeremiah M. Scharf, Carol A. Mathews, Paul M. Thompson, Dan J. Stein

## Abstract

**BACKGROUND:** There have been considerable recent advances in understanding the genetic architecture of Tourette Syndrome (TS) as well as its underlying neurocircuitry. However, the mechanisms by which genetic variations that increase risk for TS - and its main symptom dimensions - influence relevant brain regions are poorly understood. Here we undertook a genome-wide investigation of the overlap between TS genetic risk and genetic influences on the volume of specific subcortical brain structures that have been implicated in TS.

**METHODS:** We obtained summary statistics for the most recent TS genome-wide association study (GWAS) from the TS Psychiatric Genomics Consortium Working Group (4,644 cases and 8,695 controls) and GWAS of subcortical volumes from the ENIGMA consortium (30,717 individuals). We also undertook analyses using GWAS summary statistics of key symptom factors in TS, namely social disinhibition and symmetry behaviour. SNP Effect Concordance Analysis (SECA) was used to examine genetic pleiotropy - the same SNP affecting two traits - and concordance - the agreement in SNP effect directions across these two traits. In addition, a conditional false discovery rate (FDR) analysis was performed, conditioning the TS risk variants on each of the seven subcortical and the intracranial brain volume GWAS. Linkage Disequilibrium Score Regression (LDSR) was used as validation of SECA.

**RESULTS:** SECA revealed significant pleiotropy between TS and putaminal (*p*=2×10^−4^) and caudal (*p*=4×10^−4^) volumes, independent of direction of effect, and significant concordance between TS and lower thalamic volume (*p*=1×10^−3^). LDSR lent additional support for the association between TS and thalamic volume (*p*=5.85×10^−2^). Furthermore, SECA revealed significant evidence of concordance between the social disinhibition symptom dimension and lower thalamic volume (*p*=1×10^−3^), as well as concordance between symmetry behaviour and greater putaminal volume (*p*=7×10^−4^). Conditional FDR analysis further revealed novel variants significantly associated with TS (*p*<8×10^−7^) when conditioning on intracranial (rs2708146, *q*=0.046; and rs72853320, *q*=0.035 and hippocampal (rs1922786, *q*=0.001 volumes respectively.

**CONCLUSION:** These data indicate concordance for genetic variations involved in disorder risk and subcortical brain volumes in TS. Further work with larger samples is needed to fully delineate the genetic architecture of these disorders and their underlying neurocircuitry.

## 1. INTRODUCTION

Tourette’s Syndrome (TS) has a global prevalence of approximately 0.85-1% (Robertson, 2015) and is characterised by repetitive motor and phonic tics, with onset before the age of 18 years (American Psychiatric Association, 2013). TS has one of the highest heritability estimates for neuropsychiatric disorders (70-80%)(Mataix-Cols et al., 2015), with 50-60% of this heritability directly attributable to single nucleotide polymorphisms (SNPs) (Davis et al., 2013). In recent years there have been significant advances in understanding the genetic architecture of TS, and in delineating other aspects of its underlying neurobiology, including its specific neuroanatomy (Robertson, 2015).

The Psychiatric Genomics Consortium Tourette Syndrome working group (PGC-TS) undertook the first genome-wide association study (GWAS) of TS, comprising 1285 cases and 4964 controls (Scharf et al., 2013). While no SNP reached genome-wide significance, the top ranking variants were enriched for genes that affect gene expression and methylation levels in the fronto-striatal circuitry, consistent with contemporary models of TS (Robertson et al., 2017). Another study used the most highly associated variants from the PGC-TS GWAS to to nominally predict TS status (*p*=0.042) in an independent cohort (609 cases and 610 controls) and accounted for 0.52% of the variance observed between cases and controls (Paschou et al., 2014). The lack of genome-wide significance at this sample size likely reflects the polygenic and heterogeneous nature of TS (Grados 2009; Cheung et al., 2007; Jankovic et al., 2010), which is further complicated by comorbidity with other psychiatric disorders, such as obsessive-compulsive disorder (OCD), autism spectrum disorders (ASD) and attention-deficit / hyperactivity disorder (ADHD) (Hirschtritt et al., 2015).

Several studies have attempted to clarify the complex nature of TS by identifying more homogenous endophenotypes and symptom dimensions (Hirschtritt et al., 2016; de Haan et al., 2015; Cavanna et al., 2011; Robertson et al., 2008). While these studies identified several classes of Tourette-related endophenotypes using multivariate methods, all were based on relatively small samples (n<1000) (de Haan et al., 2015; Cavanna et al., 2011; Robertson et al., 2008). A recent and considerably larger analysis of individuals with TS and their family members that assessed not only TS, but also OCD and ADHD (n_total_=3494), identified two cross-disorder symptom dimensions, namely social disinhibition and symmetry behaviour (Darrow et al., 2017). Social disinhibition includes uttering syllables / words, echolalia / palilalia, coprolalia / copropraxia, and obsessive urges to offend / mutilate / be destructive. Symmetry behaviour includes symmetry, evening up, checking obsessions; ordering, arranging, counting, writing-rewriting compulsions and repetitive writing tics. Social disinhibition (h2=0.35±0.03) was associated with OCD polygenic risk scores (PRS; *p*=0.02) and less strongly with TS and ADHD PRS, which did not meet statistical significance (*p*=0.11 and *p*=0.10, respectively). In contrast, symmetry behaviour (h^2^=0.39±0.03) was significantly correlated with TS PRS (*p*=0.02) and not with OCD and ADHD (*p*=0.18 and *p*=0.26, respectively).

There have also been noteworthy advances in understanding the neuronal circuitry of TS. The role of the cortico-striatal-thalamo-cortical circuits (CSTC) in TS has been emphasized (Felling and Singer, 2011), although data on changes in the volume and function of specific brain regions in the CSTC in individuals with TS is less consistent. Lower bilateral nucleus caudate volumes (Peterson et al., 2003; Bloch et al., 2006), inferior occipital volumes (Peterson et al., 2001), prefrontal cortex volumes (Greene et al., 2016; Peterson et al., 2001), corpus callosum volumes and decreased white matter connectivity (Plessen et al., 2004, 2006) have been observed in children with TS. Greater grey matter volumes have also been observed in the thalamus (Greene et al., 2016; Miller et al., 2010), hypothalamus and midbrain among children and adults with TS (Greene et al., 2016; Peterson et al., 2007). Amygdalar volume has been reported to be greater in children and lower in adulthood (Peterson et al., 2007).

Little work to date has, however, focused on pleiotropy or concordance of genetic risk for TS, and genetic variants that influence subcortical brain volume. Pleiotropy refers to a SNP that affects both phenotypes, regardless of whether the effect direction is the same for both. Concordance, however, requires that the SNP has the same direction of effect for both phenotypes. The Enhancing Neuroimaging Genetics through Meta-analysis (ENIGMA) consortium recently performed a GWAS of structural brain MRI scans of 30,717 individuals (Hibar et al., 2015). This study identified novel genetic variants associated with the volumes of the putamen and caudate nucleus (Hibar et al., 2015) and subsequently detected their overlap with OCD risk variants (Hibar et al., 2018). ENIGMA provides an opportunity to examine the relationship between GWAS data in TS with the genetic contributions to regional brain volumes. Here we aim to assess genetic concordance for TS and specific symptom profiles (i.e., social disinhibition and symmetry behaviour) with the volume of relevant subcortical and intracranial brain regions. We used summary statistics from the ENIGMA subcortical and intracranial brain volumes GWAS (Hibar et al., 2015) and the most recent PGC-TS GWAS (Scharf et al., 2013 and unpublished data).

## 2. METHODS

### 2.1 Description of original association studies

We obtained summary statistics of adult European ancestry participants (4,644 cases and 8,695 controls) and 9,076,550 SNPs from the most recent PGC-TS GWAS (Scharf et al., 2013 and unpublished data). Approximately half of this cohort also had either comorbid OCD or ADHD. A subset of cases had information regarding symmetry behaviour (n= 1,419) and social disinhibition (n= 1,414) symptom classes. Participants were diagnosed using the DSM-IV-TR (American Psychiatric Association, 2000) by trained clinicians. Additionally, we used GWAS summary statistics from the ENIGMA Consortium meta-analysis of subcortical brain volumes across 50 cohorts including MRI scans of 30,717 individuals and 9,702,043 SNPs (Hibar et al., 2015). This cohort consists of healthy controls (79%) as well as patients (21%) diagnosed with neuropsychiatric disorders (including anxiety, Alzheimer’s disease, ADHD, major depression, bipolar disorder, epilepsy and schizophrenia). A direct comparison of the GWAS summary statistics between the full ENIGMA results (including patients) and a subset of ENIGMA results (excluding patients) showed that they were very highly correlated (Pearson’s r > 0.99) for all brain traits (Hibar et al., 2015) Prior to the analyses here, we verified that there was no overlap between the cohorts included in the TS and brain volume GWASs. The brain volume GWAS data is comprised of separate GWASs of seven subcortical brain volumes (nucleus accumbens, amygdala, caudate nucleus, hippocampus, globus pallidus, putamen, thalamus) and total intracranial volume. GWAS test statistics were genome-controlled to adjust for spurious inflation factors. All cohort studies were approved by a local ethics board prior to subject recruitment and all subjects gave written, informed consent before participating.

### 2.2 Post-processing of genetic data

To statistically compare the TS and brain volume GWASs, we used the 7,682,991 SNPs that passed quality control and filtering rules in all datasets. With these data, we performed a clumping procedure in PLINK (Purcell et al., 2007) to identify an independent SNP from every linkage disequilibrium (LD) block across the genome. The clumping procedure was performed separately for each of the 8 brain volume GWASs using a 500 Kb window, with SNPs in LD (r^2^ > 0.2), in the European reference samples from the 1000 Genome Project (Phase 1, version 3). The SNP with the lowest p-value within each LD block was selected as the index SNP representing that LD block and all other SNPs in the LD block were dropped from the analysis. The result after applying the clumping procedure was a total of 8 independent sets of SNPs representing the total variation explained across the genome conditioned on the significance in each brain volume GWAS. For each of these 8 sets of SNPs, we then determined the corresponding TS GWAS test statistic for each independent index SNP and used these datasets for the subsequent analyses.

### 2.3 Tests of pleiotropy and concordance

We used SNP Effect Concordance Analysis (SECA) (see Nyholt et al., 2014 for details of the SECA analysis) to test for genetic pleiotropy - the same SNP affecting two traits - and concordance - the agreement in SNP effect directions across these two traits - between TS, social disinhibition or symmetry behaviour and all 7 subcortical structures and ICV. Given the number of tests performed, we set a Bonferroni-corrected significance level at P* = 0.05/48 = 1,042×10^−3^ to account for the 3 traits tested (TS, social disinhibition and symmetry behaviour), the 8 brain volumes and the two tests performed (SECA and LDSR).

### 2.4 Conditional false discovery rate to detect TS, social disinhibition or symmetry behaviour risk variants

We also examined if conditioning the results of the TS GWAS on genetic variants that influence brain volume could improve our ability to detect variants associated with TS (Sun et al., 2006; Andreassen et al., 2013). At this sample size the analyses were underpowered to investigate the symptom dimensions separately. For a given brain volume phenotype, we selected subsets of SNPs at 14 false discovery rate (FDR) thresholds *q-values* ≤ (1×10^−5^, 1×10^−4^, 1×10^−3^, 0.01, 0.1, 0.2, 0.3, 0.4, 0.5, 0.6, 0.7, 0.8, 0.9, 1) and looked up the corresponding p-values for each SNP subset in the TS GWAS and the social disinhibition and symmetry behaviour symptom clusters. Next we applied the FDR method (Benjamini and Hochberg, 1995) to each subset of p-values in the TS GWAS and the social disinhibition and symmetry behaviour symptom clusters. Individual SNPs were considered significant if the p-value was lower than the significance threshold allowing for a False Discovery Rate (FDR) of 5% conditioned on any subset of SNPs from the brain volume GWASs. The LD-pruned data are still required for the conditional FDR SNP analysis because regions with varying amounts of SNPs within an LD block can affect the ranking and re-ranking of SNPs under the conditional models. However, the chosen SNP included in the model is likely just a “proxy” for SNPs in the LD block and should not necessarily be considered a causal variant or even the most significant SNP in terms of its overlap between traits.

### 2.5 Estimating genetic correlation using LD score regression

In order to replicate significant findings from our primary analysis with SECA, we used an alternative method, LD score regression (LDSR), which estimates a genetic correlation between two trait pairs based on the GWAS summary statistics of each trait analysed separately (Bulik-Sullivan et al., 2015a, 2015b). LDSR estimates a genetic correlation with a fitted linear model of Z-scores obtained from the product of significance statistics for each SNP in a given set of GWAS results compared to the level of linkage disequilibrium at a given SNP. SNPs in high LD are expected to have high Z-scores in polygenic traits with common genetic overlap (Bulik-Sullivan et al., 2015a). We used the ldsc program (https://github.com/bulik/ldsc) to perform LDSR following the methods outlined in Bulik-Sullivan et al., 2015a. LDSR was underpowered at this sample size to analyse the amygdala, as well as the TS symptom clusters. Therefore, LDSR was only used to test for an association with the broader TS phenotype and the remaining 7 brain regions of interest.

## RESULTS

### 3.1 Evidence for pleiotropy between brain volume and TS and related symptom cluster risk variants

We found significant evidence of global pleiotropy - same SNP, regardless of effect direction-between variants that infer risk for the broader TS phenotype and variants that are associated with lower putamen volumes (**Table 1**, p=2×10^−4^) and caudal volumes (**Table 1**, p=4×10^−4^). Further, we found nominally significant (*p*<0.05) evidence of pleiotropy between TS risk variants and variants associated with lower ICV, accumbens, and thalamus volumes and greater hippocampus volume (**Table 1**). No evidence for global pleiotropy was found for either the social disinhibition (**Table 2**) or symmetry behaviour (**Table 3**) TS symptom clusters.

**Table 1:** SECA results for TS whole cohort.

| Trait 1 | Trait 2 | P-value Pleiotropy | CI Pleiotropy | P-value Concordance | CI Concordance | Direction |
| --- | --- | --- | --- | --- | --- | --- |
| TS | Intracranial volume | 0.001** | 0.001-0.002 | 0.022** | 0.019-0.025 | - |
|  | Accumbens | 0.007** | 0.006-0.009 | 0.054* | 0.049-0.058 | - |
|  | Amygdala | 1 | 1-1 | 0.048** | 0.044-0.053 | + |
| | Caudate | $4 \times 10^{-4}$*** | $1 \times 10^{-4}$-0.001 | 0.016** | 0.013-0.018 | - |
|  | Hippocampus | 0.009** | 0.007-0.011 | 0.07* | 0.065-0.075 | + |
|  | Pallidum | 0.006** | 0.004-0.007 | 0.257 | 0.249-0.266 | - |
| | Putamen | $2 \times 10^{-4}$*** | $5.48 \times 10^{-5}$-0.007 | 1 | 1-1 | - |
| | Thalamus | 0.009** | 0.008-0.012 | $3 \times 10^{-4}$*** | $1 \times 10^{-4}$-0.001 | - |

TS= Tourette's syndrome, CI= confidence interval, Bonferroni corrected $p$ -value = $0.05/48 = 1.042 \times 10^{-3}$ ,
\*\*\*= significant, \*\* = nominally significant ( $p < 0.05$ ), \* = trending significance ( $p < 0.1$ )

**Table 2:**
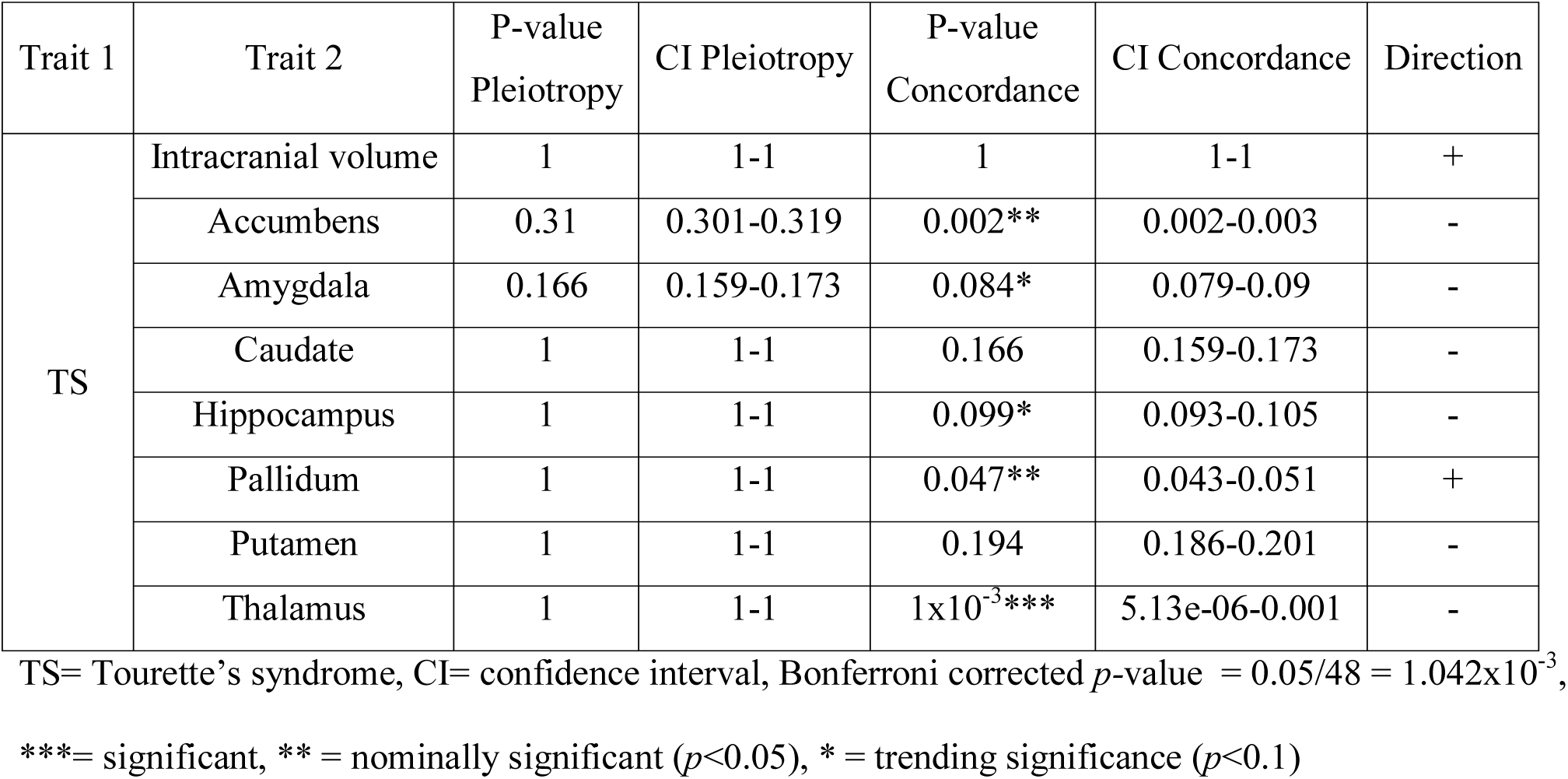
SECA results for TS social disinhibition endophenotype.

**Table 3:** SECA results for TS symmetry endophenotype.

| Trait 1 | Trait 2 | P-value<br>Pleiotropy | CI Pleiotropy | P-value<br>Concordance | CI Concordance | Direction |
| --- | --- | --- | --- | --- | --- | --- |
| TS | Intracranial volume | 1 | 1-1 | 0.191 | 0.183-0.198 | - |
|  | Accumbens | 0.304 | 0.296-0.314 | 0.263 | 0.255-0.272 | - |
|  | Amygdala | 0.298 | 0.289-0.307 | 0.002** | 0.001-0.003 | - |
|  | Caudate | 0.304 | 0.295-0.313 | 0.017** | 0.015-0.02 | + |
|  | Hippocampus | 1 | 1-1 | 0.07* | 0.065-0.075 | + |
|  | Pallidum | 1 | 1-1 | 0.417 | 0.408-0.427 | + |
|  | Putamen | 0.313 | 0.304-0.322 | 0.001*** | 0-0.001 | + |
|  | Thalamus | 1 | 1-1 | 1 | 1-1 | - |

TS= Tourette's syndrome, CI= confidence interval, Bonferroni corrected $p$ -value = $0.05/48 = 1.042 \times 10^{-3}$ ,
\*\*\*= significant, \*\* = nominally significant ( $p < 0.05$ ), \* = trending significance ( $p < 0.1$ )

### 3.2 Evidence for concordance between brain volumes and TS and related symptom clusters risk variants

We found significant evidence of negative concordance - same SNP, same direction of effect - between TS and thalamic volume (**Table 1**, *p*=3×10^−4^), indicating an association between TS genetic risk and lower thalamic volume. Further, we found nominally significant (*p*<0.05) negative concordance between the ICV and caudate with TS (**Table 1**). Nominally significant positive concordance was found between the amygdala and TS. Significant negative concordance was also identified between the social disinhibition behaviour symptom cluster and the thalamus (**Table 2**, *p*=1×10^−3^) as well as marginally significant negative concordance with the accumbens (*p*=2.3×10^−3^) and positive concordance with the pallidum (*p*=0.47×10^−2^). Significant positive concordance was also identified between symmetry behaviour and putamen volume (**Table 3**, *p*=7×10^−4^). Further, evidence for marginally significant negative concordance between this symptom cluster and the amygdala (*p*=2×10^−3^) and positive concordance with the caudate (*p*=1.7×10^−2^) were identified.

### 3.4 Replication of subcortical brain volumes and TS genetic risk overlap using LDSC

Replication using LDSR lent trending support for an association between TS risk and the thalamus (**Table 4**, *p*=5.85×10^−2^). No association between TS and pallidum volume was observed using this technique.

**Table 4:** LDSC results for TS whole cohort and brain volume overlap.

| Trait 1 | Trait 2 | rg | SE | 95% CI | P-value |
| --- | --- | --- | --- | --- | --- |
| TS | Intracranial volume | -0.122 | 0.075 | -0.269 - 0.025 | 0.106 |
|  | Accumbens | -0.428 | 0.582 | -1.568 - 0.712 | 0.462 |
|  | Amygdala | - | - | - | - |
|  | Caudate | -0.037 | 0.087 | -0.208 - 0.134 | 0.672 |
|  | Hippocampus | -0.015 | 0.097 | -0.205 - 0.175 | 0.878 |
|  | Pallidum | -0.002 | 0.106 | -0.21 - 0.206 | 0.983 |
|  | Putamen | -0.023 | 0.091 | -0.201 - 0.155 | 0.801 |
|  | Thalamus | -0.227 | 0.12 | -0.462 - 0.008 | 0.059** |

TS= Tourette's syndrome, CI= confidence interval, Bonferroni corrected $p$ -value = $0.05/48 = 1.042 \times 10^{-3}$ ,
\*\*\*= significant, \*\* = nominally significant ( $p < 0.05$ ), \* = trending significance ( $p < 0.1$ )

### 3.3 Genetic variants influencing subcortical brain volumes provide improved ability to detect TS risk variants

We performed a conditional FDR analysis, conditioning the TS risk variants on each of the 8 brain volume GWAS (**Table 5**). When conditioning the TS analysis on variants that influence ICV, rs2708146 (*q*=0.046) and rs72853320 (*q*=0.035) were significantly associated with both traits. Conditioning on the hippocampus also revealed an association between rs1922786 (*q*=0.001). No significant associations (*q*<0.05) were identified when conditioning TS on the other 6 brain GWAS.

**Table 5:** Conditional FDR results for TS and endophenotypes with brain volume overlap.

| Structure | SNP | Chr | BP | EA | NEA | Freq | Nearest gene | Distance to Gene (bp) | Effect in Brain ± Standard Error | P-value in Brain | Effect in TS ± Standard Error | P-value in TS | q-value |
| --- | --- | --- | --- | --- | --- | --- | --- | --- | --- | --- | --- | --- | --- |
| Intracranial volume | rs2708146 | 2 | 58955953 | A | G | 0,5428 | LINC01122 | intronic | 484.913± 1901.256 | 0.799 | 0.141± 0.027 | 1.3x10<sup>-7</sup> | 0.046 |
| | rs72853320 | 6 | 36623338 | A | G | 0,1331 | RNU1-88P | 15889 | -3769.398± 2836.113 | 0.184 | 0.205± 0.04 | 2.42x10<sup>-7</sup> | 0.035 |
| Hippocampus | rs1922786 | 2 | 58863573 | A | G | 0,6556 | LINC01122 | intronic | -17.527± 4.895 | 0 | 0.14± 0.028 | 7.93x10<sup>-7</sup> | 0.001 |

The chromosome (Chr) and base pair (BP) are given in h19b37 coordinates. The Effect in Brain and Effect in TS (Tourette's syndrome) are both given in terms of the effect allele (EA). The non-effect allele (NEA) is also shown. The allele frequency (Freq) corresponds to the effect allele. Tagging SNP corresponds to the most significant variant in a given LD block (if different from the SNP chosen based on clumping in the brain volume GWAS). LINC01122 = long intergenic non-protein coding RNA 1122, RNU1-88P = RNA, U1 small nuclear 88, pseudogene

## DISCUSSION

Of the eight brain traits investigated using SECA, associations were found between genetic risk for TS and for lower thalamic, putaminal and caudal volumes. These analyses of the clinical TS phenotype were further supported by SECA findings using symptom-based phenotypes; there was a significant association between genetic risk for lower thalamic volume and for the social disinhibition symptom cluster as well as a significant association between genetic risk for greater putaminal volume and for the symmetry symptom cluster. Further, three SNPs were associated with both TS and the volumes of the hypothalamus and ICV.

Associations of genetic risk for TS with genetic risk for alterations in thalamic and striatal volumes is consistent with the emphasis of neuroanatomical models of TS on these structures (Jeffries et al., 2002; Berardelli et al., 2003). The thalamus has widespread projections affecting many aspects of cognition and motor function (Shipp, 2004), and is a target for deep brain stimulation in the treatment of refractory TS (Porta et al., 2009). While greater thalamic volumes have been reported in TS (Greene et al., 2016; Miller et al., 2010), lower thalamic volumes have been identified in paediatric patients with TS (Makki et al., 2008). The dorsal striatum is also involved in motor function (Marchand et al., 2008), as well as various types of learning (Ell et al., 2006; Seger, 2005), inhibitory control of action and reward systems (Villablanca, 2010). Evidence for both greater (Ludolph et al., 2007; Roessner et al., 2011) and lower putaminal volumes (Peterson et al., 2003; Bloch et al., 2005; Greene et al., 2016) have been reported in both children and adults with TS, with lower putaminal volumes reported in TS with comorbid OCD (Peterson et al., 2003). Lower caudal volumes in adults and children have been associated with TS (Peterson et al., 2003; Bloch et al., 2005), which are particularly apparent in individuals with comorbid ADHD (Peterson et al., 2003), and are associated with tic severity and OCD symptoms (Bloch et al., 2005).

The three variants and associated with TS after conditioning on hypothalamic (rs1922786) and ICV (rs2708146 and rs72853320) volumes have not previously been associated with a particular phenotype. rs2708146 and rs1922786 are intronic variants located within the long intergenic non-protein coding RNA 1122 (*LINC01122*) gene on chromosome 2 and rs72853320 is an intergenic variant closest to the RNA U1 small nuclear 88 pseudogene (*RNU1-88P*) on chromosome 6. To date, little is known about the function of these genes, which are expressed in a broad range of tissue types, including the brain (Ensembl: https://www.ensembl.org/index.html, accessed on 5 April 2018). Other variants within *LINC01122* have previously been associated with red blood cell count, mean corpuscular haemoglobin and mean corpuscular volume in a GWAS investigating 36 blood cell phenotypes among 173,480 participants from the UK Biobank (Astle et al., 2016). This study identified a total of 2,706 variants associated with these phenotypes, located within or near genes that are involved in pathways that have been implicated in schizophrenia, autoimmune diseases and cardiovascular disorders. (Astle et al., 2016). As little is known about the genes and variants identified in our study, additional replications and functional studies are needed.

The analyses here complement prior work on OCD, where we found significant concordance between OCD risk variants and variants that are related to greater putaminal volume (*p*=8.0 × 10^−4^) (Hibar et al., 2018). In addition, the CSTC has been implicated in both disorders (Berardelli et al., 2003; Rauch and Britton, 2010). Our previous study on OCD did, however, also identify significant concordance between greater nucleus accumbens volume and OCD and conditional FDR only revealed significant associations with OCD when conditioned on putaminal, amygdalar, and thalamic volumes (Hibar et al., 2018). Differences between OCD and TS are consistent with genetic analyses which have emphasized that despite their relatedness, these conditions have different genetic architectures (Davis et al., 2013; Yu et al., 2015). The findings here arguably support the proposal in the eleventh revision of the International Classification of Diseases (ICD-11) to classify TS both as a neurodevelopmental disorder and as an obsessive compulsive related disorder (OCRD) (Stein et al., 2016; Robertson, 2015).

Several limitations of this work should be emphasized. First, the LDSR analysis only lends trending support for the association between the thalamus and TS. LDSC was further underpowered to perform all the tests between brain volume and TS. Validation in an even larger cohort is, therefore, still necessary. Second, the analysis could be biased if overlapping participants were present in the studies contributing to the consortia. We verified that the cohorts as whole did not overlap and individual overlap is, therefore, likely to be minimal. Third, the ENIGMA GWASs of brain volumes contain cohorts with healthy controls as well as patients diagnosed with neuropsychiatric disorders (including anxiety, Alzheimer’s disease, ADHD, major depression, bipolar disorder, epilepsy and schizophrenia), which may bias the interpretation of our findings and how they relate to TS. However, a direct comparison of the GWAS summary statistics between the full ENIGMA results (including patients) and a subset of ENIGMA results (excluding patients) showed that they were very highly correlated (r^2^ > 0.99) for all brain traits (Hibar et al., 2015). This suggests that the pattern of effects in the brain volume GWAS is not likely driven by disease status. Fourth, approximately half of the TS cohort also has comorbid OCD. It is possible that stronger and more specific associations may be revealed in a cohort with TS only. Fifth, the relationship between gene variants influencing brain volume and neuropsychiatric risk may be influenced by a range of confounders, including environmental factors such as stress and medication effects, which have effects on brain volume and disease risk independent of genetics. Discovering the pathway by which gene variants influencing brain volume also create risk for TS and its symptom clusters is therefore susceptible to influence by environmental factors, which might obscure genetic relationships. However, an endeavour to find the genetic overlap between brain volume and disorder risk using the largest datasets to date is still worthwhile as it can potentially provide insights into the disorders.

In conclusion, this study implicated genetic overlap between genetic variants influencing the volumes of the thalamus, putamen and caudate in TS and its symptom dimensions. Further, this study identified three SNPs associated with TS and volumes of the hypothalamus and ICV. Indeed, these data are the first to show an overlap between risk for genes for TS and for brain circuitry. The correlations with putaminal and thalamic volumes are consistent with a broad range of previous neuroimaging work. Emerging collaborations and consortia, such as ENIGMA-TS, aim to continue to increase sample size, which will enhance statistical power in future iterations of this analysis. Additional work focusing on a range of other methodologies to assess genetic overlap may also be useful, following along the lines of recent work in schizophrenia (Lee et al., 2016; Franke et al., 2016; Mufford et al., 2017). Such studies have used partitioning-based heritability analysis (Yang et al., 2011) and conjunction analysis (Nichols et al., 2005) to identify genetic variants associated with both schizophrenia risk and altered brain volumes. These approaches may also be useful in future work on TS and its symptom dimensions.

## Acknowledgements

**PGC-TS members:** Harald Aschauer; Gil Atzmon; Cathy Barr; Csaba Barta; Robert Batterson; Fortu Benarroch; Chester Berlin; Gabriel Berrio; Julia Bohnenpoll; Lawrence Brown; Ruth Bruun; Randy Buckner; Cathy Budman; Julio Cardona Silgado; Danielle Cath; Keun-Ah Cheon; Sylvain Chouinard; Barbara Coffey; Giovanni Coppola; Nancy Cox; Sabrina Darrow; Lea Davis; Christel Depienne; Andrea Dietrich; Yves Dion; Valsamma Eapen; Lonneke Elzerman; Thomas Fernandez; Nelson Freimer; Odette Fründt; Blanca Garcia-Delgar; Donald Gilbert; Marco Grados; Erica Greenberg; Dorothy Grice; Varda Gross-Tsur; Julie Hagstrøm; Andreas Hartmann; Johannes Hebebrand; Tammy Hedderly; Gary Heiman; Luis Herrera; Isobel Heyman; Matthew Hirschtritt; Pieter Hoekstra; Hyun Ju Hong; Alden Huang; Chaim Huyser; Laura Ibanez-Gomez; Cornelia Illmann; Joseph Jankovic; Judith Kidd; Kenneth Kidd; Young Key Kim; Young-Shin Kim; Robert King; Yun-Joo Koh; Anastasios Konstantinidis; Sodahm Kook; Samuel Kuperman; Roger Kurlan; James Leckman; Paul C. Lee; Bennett Leventhal; Thomas Lowe; Andrea Ludolph; Claudia Lührs da Silva; Gholson Lyon; Marcos Madruga-Garrido; Irene Malaty; Athanasios Maras; Carol A. Mathews; William McMahon; Sandra Mesa Restrepo; Pablo Mir; Astrid Morer; Kirsten Müller-Vahl; Alexander Münchau; Tara Murphy; Allan Naarden; Peter Nagy; Benjamin Neale; Markus Noethen; William Ochoa; Michael Okun; Lisa Osiecki; Peristera Paschou; David Pauls; Christopher Pittenger; Kerstin Plessen; Yehuda Pollak; Danielle Posthuma; Eliana Ramos; Victor Reus; Renata Rizzo; Mary Robertson; Veit Roessner; Josh Roffman; Guy Rouleau; Andres Ruiz-Linares; Paul Sandor; Jeremiah Scharf; Monika Schlögelhofer; Eun-Young Shin; Harvey Singer; Jan Smit; Jordan Smoller; Dong-Ho Song; Jungeun Song; Mara Stamenkovic; Matthew State; Manfred Stuhrmann; Jae-Hoon Sul; Urszula Szymanska; Zsanett Tarnok; Jay Tischfield; Fotis Tsetsos; Jennifer Tübing; Ana Valencia Duarte; Frank Visscher; Sina Wanderer; Tomasz Wolanczyk; Martin Woods; Yulia Worbe; Dongmei Yu; Ivette Zelaya; Samuel Zinner

